# Hippocampal theta oscillations support successful associative memory formation

**DOI:** 10.1101/2020.05.26.116517

**Authors:** Srinivas Kota, Michael D. Rugg, Bradley C. Lega

## Abstract

1.

Models of memory formation posit that recollection as compared to familiarity-based memory depends critically on the hippocampus, which binds features of an event to its context. For this reason, the contrast between study items that are later recollected versus those that are recognized on the basis of familiarity should reveal electrophysiological patterns in the hippocampus selectively involved in associative memory encoding. Extensive data from studies in rodents support a model in which theta oscillations fulfill this role, but results in humans results have not been as clear. Here, we employed an associative recognition memory procedure to identify hippocampal correlates of successful associative memory encoding and retrieval in patients undergoing intracranial EEG monitoring. We identified a dissociation between 2– 5 Hz and 5–9 Hz theta oscillations, by which 2–5 Hz oscillations uniquely were linked with successful associative memory in both the anterior and posterior hippocampus. These oscillations exhibited a significant phase reset that also predicted successful associative encoding, distinguished recollected from familiar items at retrieval, and contributed to reinstatement of encoding-related patterns that distinguished these items. Our results provide direct electrophysiological evidence that 2–5 Hz hippocampal theta oscillations support the encoding and retrieval of memories based on recollection but not familiarity.

**Significance Statement:** Extensive fMRI evidence suggests that the hippocampus plays a selective role in recollection rather than familiarity, during both encoding and retrieval. However, there is little or no electrophysiological evidence that speaks to whether the hippocampus is selectively involved in recollection. Here, we used intracranial EEG from human participants engaged in an associative recognition paradigm. The findings suggest that oscillatory power and phase reset in the hippocampus are selectively associated with recollection rather than familiarity-based memory judgements. Furthermore, reinstatement of oscillatory patterns in the hippocampus was stronger for successful recollection than familiarity. Collectively, the findings support a role for hippocampal theta oscillations in human episodic memory.

## 3. Introduction

Models of episodic memory encoding posit that hippocampal theta oscillations fulfill the critical role of organizing neuronal ensembles to link together item and contextual representations (Eichenbaum, 2017; Staudigl & Hanslmayr, 2013). Numerous experimental findings have established the importance of theta oscillations for associative memory using spatial paradigms in rodents, including the observation that interventions that disrupt theta oscillations attenuate associative encoding (Stella & Treves, 2011; Yamaguchi et al., 2007). However, the role of hippocampal theta oscillations in *human* associative memory has proven more difficult to establish. In spatial navigation and free recall, theta power increases in the 2-5 Hz frequency range have been reported to predict successful memory encoding, and additional properties such as phase amplitude coupling, phase reset and phase synchronization have also been described for oscillations in this frequency range (Ekstrom et al., 2005; Lega, Jacobs, & Kahana, 2012; Lin et al., 2017). However, most studies examining hippocampal oscillatory changes during mnemonic processing report power decreases in the 2–10 Hz band (Fellner et al., 2016; Greenberg, Burke, Haque, Kahana, & Zaghloul, 2015). By contrast, intracranial EEG (iEEG) studies employing recognition memory paradigms report properties such as phase amplitude coupling, phase reset, and phase locking of single units that include faster theta frequencies up to 9 Hz (Axmacher, Mormann, Fernández, Elger, & Fell, 2006; Fell & Axmacher, 2011). On the one hand, these findings support the general importance of theta oscillations in human memory, but observing these properties within a simple recognition (rather than an episodic or spatial navigation paradigm) may not necessarily support a model by which theta oscillations specifically support item-context associations, as predicted from rodent data.

This question may in turn be related to possible differences between 2–5 Hz ‘slow’ theta oscillations and 5–9 Hz ‘fast’ theta oscillations in human iEEG recordings from the hippocampus (Jacobs, 2014). While many reports of hippocampal theta do not attempt to segregate oscillations within the broad 2–9 Hz frequency range, human data suggest that it is oscillations below 5 Hz that exhibit mnemonic correlates consistent with predictions from rodent experiments, including power increases during successful as compared to unsuccessful encoding and phase amplitude coupling (Lega, Burke, Jacobs, & Kahana, 2014; Lega et al., 2012; Lin et al., 2017). In the present study, we sought to clarify the role of hippocampal theta oscillations by testing directly whether oscillations in ‘low’ and ‘high’ frequency ranges distinguish successful versus unsuccessful associative memory encoding and retrieval.

A central tenet of episodic memory retrieval is the concept of reinstatement, meaning that the brain patterns present at the time of encoding recur in the brain at the time of retrieval. Across both invasive and noninvasive modalities, evidence of memory reinstatement has been demonstrated in human studies (Thakral, Wang, & Rugg, 2015; Yaffe et al., 2014). A clear prediction of these data related to the associative recognition task is that reinstatement (as quantified by distance metrics such as cosine similarity) should be greater for recollected items (associative hits) as compared to items recalled on the basis of familiarity (associative misses). To our knowledge, this has not been previously tested using hippocampal recordings.

We employed an associative recognition procedure previously validated in human fMRI studies (de Chastelaine, Mattson, Wang, Donley, & Rugg, 2016a, 2016b). In this procedure, participants first study a series of item pairs. In the subsequent retrieval task, test items compromise a mix of ‘intact’ pairs (re-presentations of studied pairs), ‘rearranged’ pairs (pairs comprising words presented on different study trials) and ‘new’ pairs (comprising previously unstudied items). Test pairs that are correctly recognized as intact (termed associative hits) are deemed to have been successfully recollected, while intact pairs incorrectly classified as rearranged signify failed recollection but intact familiarity (see (de Chastelaine et al., 2016a, 2016b) for a detailed rationale of these operationalizations). A key feature of this procedure is that it permits a direct contrast between patterns of brain activity linked with successful versus unsuccessful associative recollection, making it well-suited for neurophysiological examinations of hippocampal processing supporting associative memory. Our primary goal was to ascertain whether, during encoding, theta power would differ according to whether a study item went on to be successfully recollected (associative hit) or to be wrongly judged as rearranged (associative miss). Thus, the contrast between associative hits versus misses is akin to that between recollection- and familiarity–based memory retrieval, and we use these terms synonymously in describing our results.

Of importance, we employed as participants twenty individuals in whom intracranial electrodes had been inserted into both anterior and posterior hippocampus. This allowed us to interpret our findings in light of models of hippocampal longitudinal specialization (Poppenk, Evensmoen, Moscovitch, & Nadel, 2013). The associative recognition paradigm further permitted us to examine differences in oscillations elicited by associative hits and misses (recollection versus familiarity-based memory) both at the time of encoding and retrieval. This is highly pertinent since fMRI BOLD data indicates that hippocampal activity preferentially distinguishes these item type at both encoding and retrieval (de Chastelaine et al., 2016a).

## 4. Materials and Methods

Twenty adult participants (22-54 years; mean age = 34 years, SD = 10 yrs; 10 female; 16 right-handed, table 1) with pharmacologically resistant epilepsy were implanted with stereotactic EEG electrodes to detect seizure location. The number and location of the intracerebral electrodes implanted in the patients were determined exclusively based on the clinical need. The participants were recruited from the Epilepsy Monitoring Unit at the University of Texas, Southwestern Medical Center, Dallas. Each participant provided informed consent prior to participation in the research study in accordance with the UT Southwestern’s Institutional Review Board. Data collected from seven additional individuals were excluded due to insufficient number of trials or inadequate behavioral performance, falling below an *a priori* defined cut-off of 0.1 for associative recognition accuracy (pIntact|Intact - pIntact|Rearranged, see behavioral results below).

**Table 1:** Demographic data.

| Subject no | Sex | Age | Duration of epilepsy (y) | Hemisphere of onset | Electrodes in AH | Electrodes in PH | Location of seizure onset |
| --- | --- | --- | --- | --- | --- | --- | --- |
| 1 | F | 38 | 14 | B | 6 | 5 | Right heterotopia, Left heterotopia |
| 2 | F | 31 | 6 | L | 1 | 2 | Left insula |
| 3 | M | 38 | 35 | L | 1 | 0 | Left orbito-frontal cortex |
| 4 | F | 22 | 9 | L | 6 | 2 | Left fusiform gyrus |
| 5 | F | 36 | 19 | R | 0 | 2 | Right anterior cingulate/right SFG prefrontal, right anterior insula/frontal operculum |
| 6 | M | 27 | 21 | B | 5 | 2 | Bilateral hippocampi |
| 7 | M | 41 | 30 | B | 4 | 0 | Amygdala, right insula |
| 8 | M | 54 | 30 | R | 1 | 2 | STS/MTG posterior |
| 9 | M | 26 | 2 | R | 5 | 2 | Right temporal lobe |
| 10 | M | 22 | 3 | B | 3 | 3 | Bilateral temporal lobe |
| 11 | F | 42 | 8 | L | 3 | 1 | Left posterior insula |
| 12 | M | 31 | 27 | R | 3 | 0 | Right hippocampus |
| 13 | F | 54 | 4 | L | 3 | 2 | Left entorhinal |
| 14 | M | 20 | 4 | B | 5 | 4 | Right collateral sulcus/fusiform gyrus/ heterotopia |
| 15 | F | 37 | 4 | L | 2 | 1 | Left temporal pole |
| 16 | F | 31 | 10 | L | 6 | 6 | Left anterior hippocampus, amygdala |
| 17 | F | 22 | 17 | L | 4 | 2 | Left hippocampus |
| 18 | M | 23 | 6 | R | 6 | 0 | Right pre-frontal/pre-motor |
| 19 | M | 45 | 12 | R | 5 | 3 | Right hippocampus |
| 20 | F | 45 | 23 | L | 6 | 2 | Left schizencephalic |

LH/RH, left/right hemisphere; B, bilateral; AH, anterior hippocampus; PH, posterior hippocampus

### 4.1. Experimental Stimuli

The experimental stimuli consisted of 320 visually presented, semantically unrelated word pairs. The words were concrete nouns ranging from 3 to 9 letters, selected from word association norms (Nelson, McEvoy, & Schreiber, 2004). 240 word pairs were presented at the time of study (encoding), with the remainder (80) held as ‘novel’ pairs presented at test (retrieval). Of the 240 word pairs seen at study, 160 were presented at test identically to how they were shown at study (intact pairs), and 80 were ‘rearranged,’ meaning the individual words had appeared previously but were presented at test paired with a different word. These were divided into two lists (120 pairs at presented at encoding, 160 presented at test). Participants completed a practice study list consisting of a separate group of 45-word pairs shown at study, and the test list for this practice list included 60-word pairs (30 intact, 15 rearranged, 15 novel).

### 4.2. Procedure

Participants were provided instructions for the both the study (encoding) task and the memory test (retrieval) prior to undertaking the experiment proper. Upon completion of the practice session, the study and test phases of the experiment were administered. All participants completed at list one study/test cycle following practice (minimum of 120 word pairs at study, 160 at test). Fig. 1 shows a schematic overview of the study and test procedures. For the study portion, participants were required to indicate with a button press which of the two objects denoted by the words was more likely to fit into the other object. After completing the study task, participants engaged in a math distractor task for 30 seconds. The task was in the form of A+B+C = ??, in which A, B, and C are single digits. There then followed a five-minute break, after which participants undertook the associative recognition test. For the memory test, participants were required to press one of the three keys to indicate whether a test pair was intact, rearranged or new. ‘Intact’ responses indicated participants recognized both words and had a specific memory of the two words being presented together at study; ‘rearranged’ responses indicated that participants recognized both words from study but had no memory of the words having previously been presented together. If participants could not recognize one or both of the words in a test pair, they made a ‘new’ response. The following terms are used throughout the paper to describe behavioral responses: ‘associative hits’ to denote correctly endorsed intact items, and ‘associative misses’ to denote intact items incorrectly identified as rearranged. Study and test instructions emphasized the need for both accuracy and speed. During both study and test phases, participants were seated comfortably in their beds. Stimuli were presented on a laptop computer that was placed on a height adjustable overbed bedside table approximately one meter in front of the participant.

**Figure 1:**
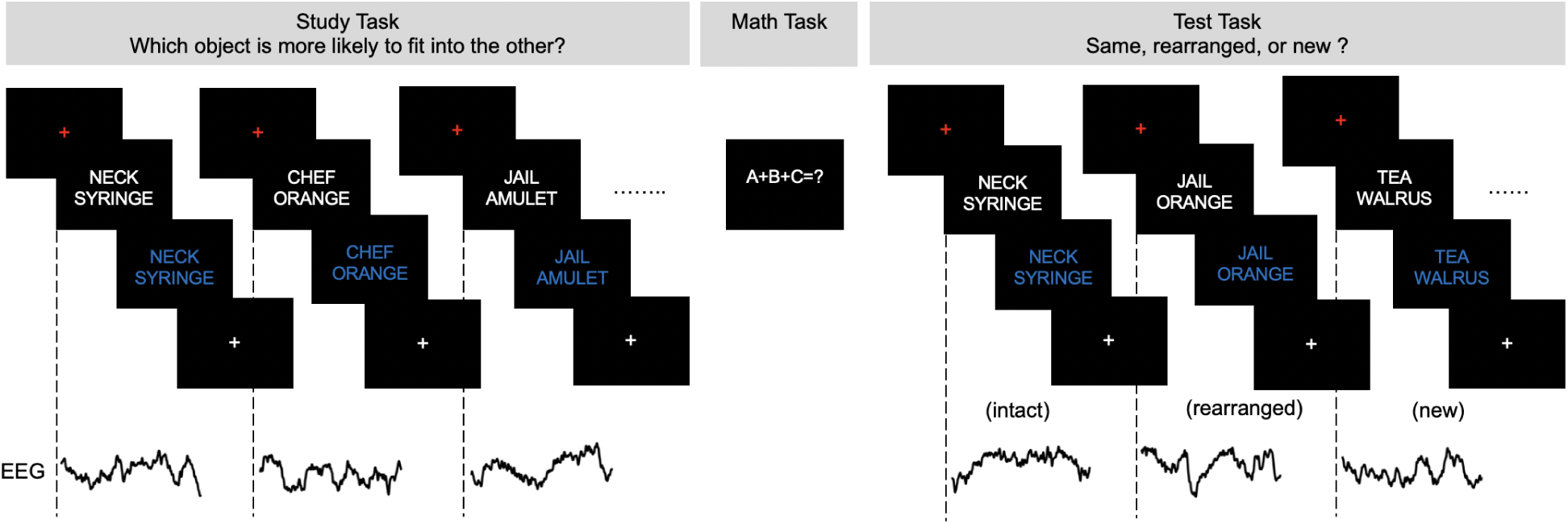
A schematic of the associative recognition paradigm showing study, distractor, and test procedures.

Study and test pairs were preceded by a red fixation cross for 0.5 s. The pairs were presented in a white uppercase Helvetica 30 font against a black background for 2 s. Then, the study pair turned from white to blue for 1 s to prompt participants’ response. This same timing was used at test. Study and test pairs were followed by a white fixation cross for 1 s (jitter 200 ms). Reaction times were measured from the onset of the word pairs on the laptop computer screen but participants were allowed to respond when font changed from white to blue.

### 4.3. EEG acquisition and preprocessing

Participants all had intracranial electrodes implanted in the anterior and posterior hippocampus (nineteen subjects with anterior electrodes, sixteen subjects with posterior electrodes, fifteen subjects contributing both left and right hemisphere hippocampal electrodes), localized by independent expert neuroradiology review, with the uncal notch set as the demarcation point between anterior and posterior regions. EEG signals from hippocampal electrodes (1-12 electrodes per hippocampus including both anterior and posterior contacts, M = 6, SD = 3) were sampled at 1 kHz on a Nihon-Kohden 2100 clinical system. EEG signals were processed offline using MATLAB software (The MathWorks, Inc., Natick, MA, US). EEG signals were re-referenced to an average of all available electrodes (across participants 116-260 electrodes, M = 189, SD = 38) after excluding noisy channels or those with frequent interictal activity based upon the report of an epileptologist. Study pairs that were correctly endorsed as intact (subsequent associative hits) and pairs that were later incorrectly identified as rearranged (subsequent associative misses) on the subsequent associative recognition test were employed in the analyses of the encoding data. Data recorded at the time of test (memory retrieval) were analyzed separately. For both study and test phases, continuous EEG signals were segmented into 1600 ms epochs time-locked to the onset of the stimulus. The EEG signals were notch filtered at 60 Hz (4 Hz bandwidth) and its harmonics. Kurtosis (ratio of fourth central moment of the data and fourth power of its standard deviation) for each EEG epoch (0 to 1600 ms) was calculated. EEG epochs that had a kurtosis value higher than four were considered to contain artifact and were excluded (Lin et al., 2017). To further eliminate artifacts, a median based artifact detection algorithm was used (Kook, Gupta, Kota, Molfese, & Lyytinen, 2008). With this approach, the median signal of the remaining EEG epochs was calculated, and a Euclidean distance from each epoch to the median signal was then computed. EEG epochs that exceeded the 95th percentile of the Euclidean distances were considered artifacts and excluded. EEG epochs were resampled to 500 Hz and power calculated for the entire epoch (0 to 1600 ms) using Morlet wavelets at 54 logarithmic frequencies ranging from 2 to 200 Hz (Lin et al., 2017).

### 4.4. Oscillatory power analysis

All analyses of oscillatory power were performed by collapsing test statistics (for example Z-scores) within a single hippocampus, such that analyses of power utilize a single data point for each subjects’ hippocampus. Subjects contributing bilateral data provided two data points. Analyses were performed separately for anterior and posterior hippocampi. To examine subsequent memory effects, power for subsequent associative hits versus subsequent associative misses was contrasted for every electrode in the anterior hippocampus and posterior hippocampus. This comparison was performed using a Wilcoxon rank-sum test by shuffling events randomly 1000 times at each time instant and frequency. A normal inverse transformation was applied to the p-value matrices of each electrode by comparing the true p value to that obtained from the shuffle distribution, converting them to Z-scores. These values were averaged for all electrodes within the hippocampus of every subject, creating a distribution of Z-scores across all hippocampi. This distribution was compared to a null hypothesis of no difference in power between associative hits versus misses using a one-sample t-test with an associated shuffle procedure (5000 shuffles). False discovery rate (Q = 0.05) was used to control the probability of Type I error. Significant results in time frequency space indicate a power difference with an FDR corrected p value < 0.05 for a minimum of one continuous cycle of an oscillation.

To further characterize slow and fast theta oscillations, power was averaged across trials for every electrode for subsequent associative hits and misses and normalized using 1 second from the prestimulus baseline period at each frequency (1 Hz steps, 2-9 Hz). Normalized power was combined for slow theta (2–5 Hz) and fast theta (5–9 Hz) for each event for all electrodes within each hippocampus. Normalized power for subsequent associative hits versus misses at every time point (2 ms separation) was compared using a Wilcoxon rank-sum test with an associated shuffle procedure (1000 shuffles). Time points with a significant difference in normalized power after FDR (Q = 0.05) correction were identified.

Averaged normalized power was calculated for time points that were significantly different for subsequent associative hits and misses for slow theta and fast theta separately for every electrode for illustrative purposes only. The above analysis approach was repeated for associative hits versus associative misses in the data from the test phase (item retrieval). Bayesian independent samples t-test (JASP version 0.11.1) was used to compare averaged normalized power for anterior and posterior hippocampus in slow theta during retrieval.

### 4.5. Oscillation Detection

To confirm that both slow and fast theta oscillations were present during successful associative memory encoding and retrieval, we used an oscillation detection algorithm. Artifact-free trials were averaged for each electrode. The averaged EEG signal was used to identify peak frequencies using the Multiple Oscillation Detection Algorithm (MODAL) for frequency ranges from 0.5 to 32 Hz (Watrous, Miller, Qasim, Fried, & Jacobs, 2018). This algorithm included a procedure to remove 1/*f* fit from the power spectrum and adaptively identify frequency bands. The Yule’Q test (Lewis-Beck, Bryman, & Liao, 2003) was employed to test whether slow and fast theta oscillations occurred simultaneously in the same hippocampal recording contact.

### 4.6. Phase Reset

We examined the phenomenon of phase reset for slow theta and fast theta. Due to a prior concern regarding heterogeneity in the precise timing and preferred phase of reset across electrodes (Rizzuto, Madsen, Bromfield, Schulze-Bonhage, & Kahana, 2006), this analysis was performed at the electrode level rather than collapsing data at the within each hippocampus (fixed effects approach). EEG signals were filtered using a Butterworth bandpass filter 2-5 Hz and 5-9 Hz. Phase values were then calculated during the study phase with the Hilbert transform at every time point (2 ms seperation) for subsequent associative hits and misses. A Rayleigh test was used to identify non-uniformity in the phase angle distribution at each time step across the epoch (0 to 1600 ms). P-values from this test were converted to Z-scores using a normal inverse transformation. This process was repeated for all electrodes in the anterior and posterior hippocampus. To test whether the phase reset values for subsequent associative hits were significantly different from those for subsequent associative misses within each electrode, a Wilcoxon rank-sum test with a 5% significance level and associated shuffle procedure was used after collapsing information for hippocampus. The resultant p-value matrices were converted to Z-scores. This procedure was also performed for associative hits and misses during the test phase. To test whether the slow theta phase reset that was observed for associative hits in the study and test phases tended to occur at similar phases, the mean phase difference between study and test was compared across electrodes using the Rayleigh test. The comparison was conducted separately for the anterior hippocampus and posterior hippocampus.

### 4.7. Reinstatement of encoding-related oscillations

Cosine similarity was used as a metric to estimate hippocampal reinstatement during successful recollection (Yaffe et al., 2014). Cosine similarity is the cosine angle between two vectors. Mathematically, it is the ratio of the dot product of two vectors and the product of Euclidean magnitudes of two vectors. It has been shown that cosine similarity and correlation methods outperform Euclidean distance for quantitating similarity between high dimensional space vectors (Huang, 2008; Strehl, Ghosh, & Mooney, 2000). Pearson’s correlation is similar to cosine similarity is applied to the mean-centered data (Van Dongen & Enright, 2012). We conducted this analysis at the electrode level (fixed effects approach). We hypothesized that cosine similarity values for successful recollection would be higher than for familiarity in both anterior and posterior hippocampus. Artifact-free trials that were common to both study and test were identified, and power was normalized at each frequency. The average power across trials was calculated at study and test separately for every electrode in the anterior hippocampus and posterior hippocampus for every frequency (54 logarithmic frequencies ranging from 2 to 200 Hz). Cosine similarity was calculated across the entire frequency spectrum (2-200 Hz), and broken down into slow theta (2-5 Hz), fast theta (5-9 Hz) and high gamma (110 to 200Hz) in a 500 ms time window with a 90% overlap (sliding window width = 50 ms) for every 500 ms in study with all 500 ms time windows at the time of test following published methods (Yaffe et al., 2014). This results in a plot of reinstatement values comparing all study (encoding) windows to all test (retrieval) windows (a reinstatement map). This was done separately at each electrode for successful recollection (associative hits) and separately for unsuccessful recollection (associative misses). The Fisher z-transformation was applied to the cosine similarity values. Z-scores were compared between successful recollection versus familiarity across electrodes using a Wilcoxon rank-sum test with an associated shuffle procedure (1000 shuffles) at each time window. Significant differences in reinstatement magnitude were those for which the p value was <0.05 following FDR correction across time windows.

## 5. Results

Twenty subjects implanted with intracranial electrodes met our minimum performance threshold while completing the AR task (see behavioral results). Further demographic details are provided in Table 1 that include subjects’ sex, age, duration of epilepsy in years, hemisphere of onset, number of electrodes in hippocampus, and seizure onset location.

### 5.1. Behavioral results

Associative recognition performance was estimated by the metric pR - the difference between the probability of an intact response to an intact test item and probability of an intact response to a rearranged item (King, de Chastelaine, Elward, Wang, & Rugg, 2015). Participants with a pR > 0.1 (probability of chance) were included in the analyses presented below. Mean (SD) of accuracy and reaction time (RT) on the study task was 0.53 (0.22) and 2515 (86) ms, respectively. The mean (SD) proportion of intact trials correctly endorsed as intact (associative hits) was 0.58 (0.17), and a mean of 0.20 (0.09) of trials were classified as associative misses. Mean (SD) pR was 0.39 (0.22). Mean reaction time (SD) for associative hits was 2608 (118) ms, and for associative misses it was 2695 (180) ms. Pair-wise *t* test revealed that the reaction time for associative misses was significantly greater than the associative hits (t19=3.20, p=0.0023).

### 5.2. Theta power distinguishes recollected from familiar items at both encoding and retrieval

Our principal hypothesis was that we would observe a difference in oscillatory power in the theta frequency range distinguishing associative hits (recollected items) from associative misses (familiar items). For our initial analysis, we did not collapse power data across frequency bands. We used a rank-sum test with an associated shuffle procedure with FDR correction to compare oscillatory power between recollected and familiar items at each time-frequency pixel separately for the study and test phases. Significant time-frequency pixels across the 2– 9 Hz theta range are shown in Figures 2A (encoding) and 3A (retrieval). During encoding, we observed significantly greater oscillatory power in both slow and fast theta frequency ranges for later recollected (subsequent associative hits) than for familiar items (subsequent associative misses) in both the anterior hippocampus (FDR corrected *p* < 0.05, effect size 0.20-0.33). In the posterior hippocampus we observed qualitatively the same pattern during study (encoding), but a fewer time-frequency pixels survived FDR correction.

**Figure 2:**
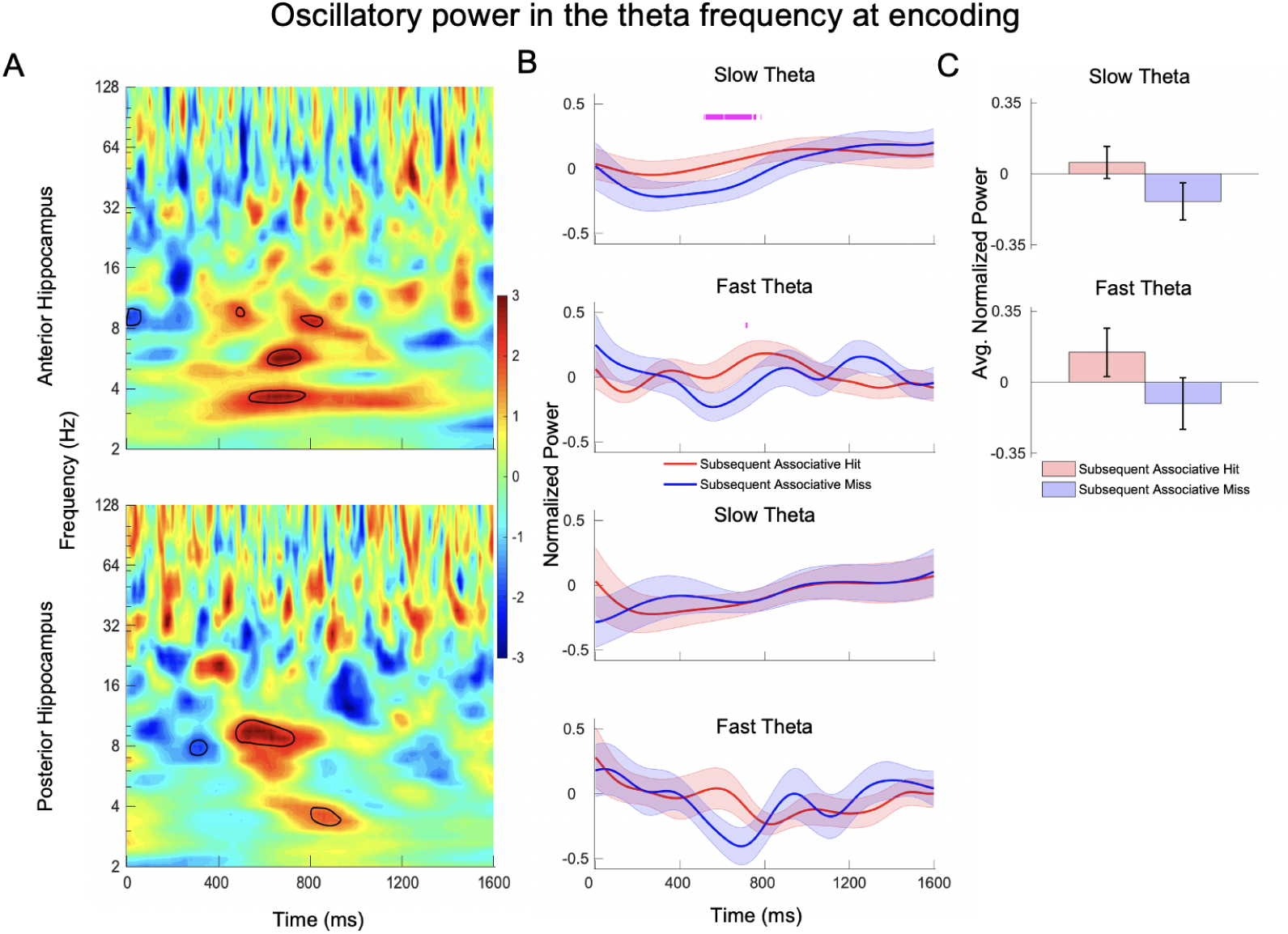
Time-frequency maps comparing power during encoding between intact items that went on to be classified at test as associative hits versus misses. A) Z-score values are plotted for anterior hippocampus (top panel) and posterior hippocampus (bottom panel). In the time-frequency plot, red indicates that the power for associative hits is significantly higher than for associative misses, and blue indicates the opposite. Time-frequency pixels in the shaded region in theta range (2-9 Hz) survived correction (FDR corrected p < 0.05, rank-sum with 5000 shuffles across subjects z-scores). B) Aggregate normalized power for slow theta and fast theta for the anterior hippocampus (top panel) and posterior hippocampus (bottom panel). Tick marks on the top in magenta indicate time points at which normalized power for subsequent associative hits is significantly higher than subsequent associative misses using rank-sum with 1000 shuffles, and cyan indicates the opposite. C) Bar plots show the averaged normalized power for the time points that are significantly different across subsequent associative hits and subsequent associative misses. Error bars indicate standard error of the mean.

To more precisely characterize differences in theta power during study, we next aggregated normalized power within each hippocampus across previously–defined 2–5 Hz and 5–9 Hz frequency ranges and contrasted power at each time point between subsequent associative hits versus subsequent associative misses. Time instants at which significant differences were detected are shown in Figure 2B (FDR corrected *p* < 0.05, effect size 0.20-0.32, rank-sum test with shuffle procedure). Results using this method were consistent with the results for the full time–frequency plots described above, We observed significantly greater slow–theta power in the anterior hippocampus for subsequent associative hits from approximately 500–800 ms but more temporally restricted effect in the fast–theta band centered around 700 ms after item presentation. We observed no time points that survived FDR correction in the posterior hippocampus.

During retrieval, oscillatory differences between associative hits and misses (and between frequency bands) were starker. Fig 3B shows that associative hits elicited power increases in the slow–theta band that were centered around 1000 ms post-stimulus onset (FDR corrected *p* < 0.05, rank-sum test) in the anterior hippocampus, consistent with results in the time– frequency plots illustrated in Figure 3A. Interestingly, in the fast–theta frequency range, a power increase *in the opposite direction* occurred around 480 ms after item onset, with greater fast– theta oscillatory power for associative misses. The same pattern was evident in the posterior hippocampus, although fewer time points survived FDR correction both for the slow and fast theta power differences (Fig 3B).

**Figure 3:**
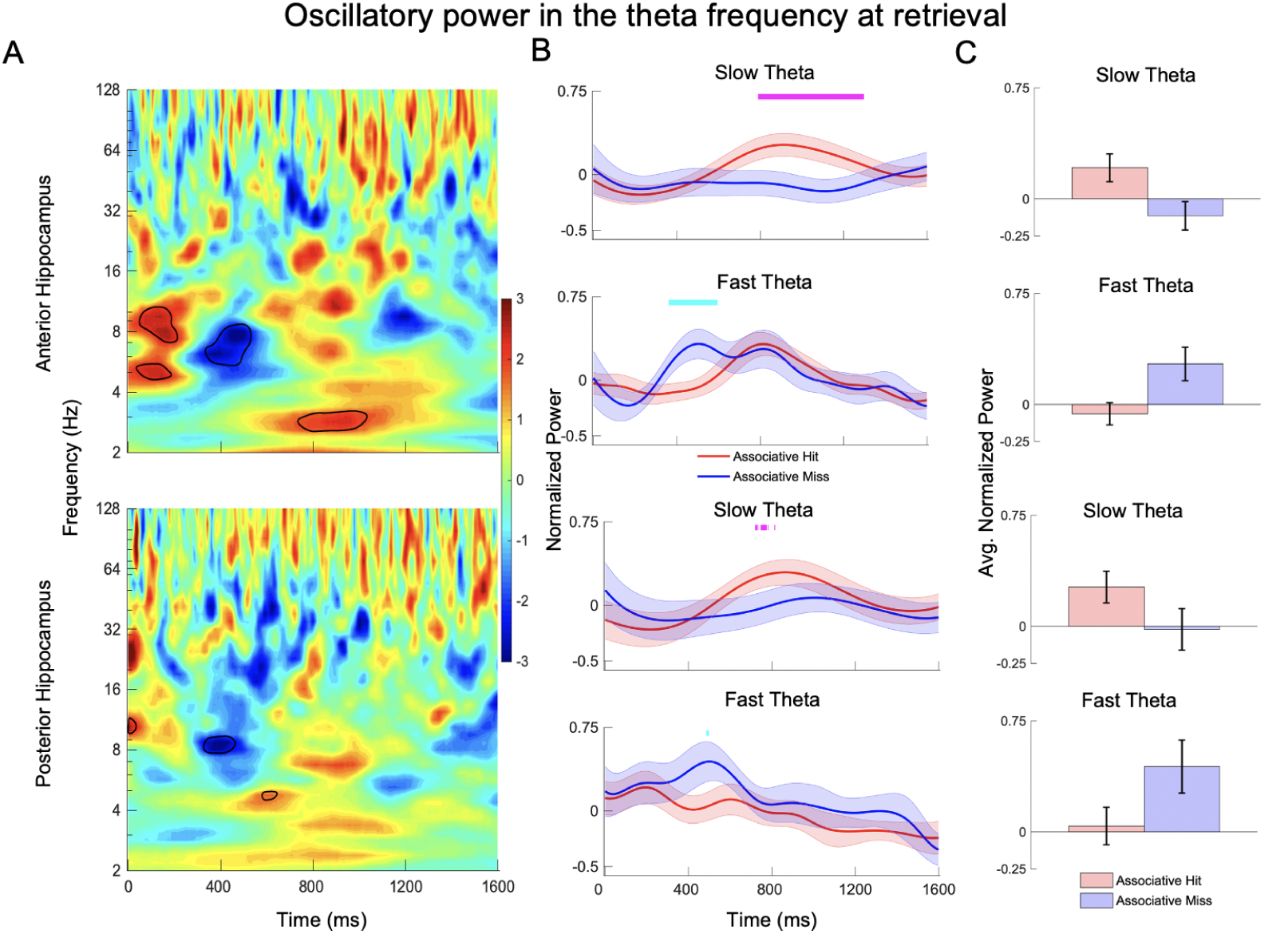
Time-frequency maps comparing power during retrieval between associative hits versus misses. A) Z-score values are plotted for anterior hippocampus (top panel) and posterior hippocampus (bottom panel). In the time-frequency plot, red indicates the power for associative hits is significantly higher than associative misses, and blue indicates the opposite. Time-frequency pixels in the shaded region in theta range (2-9 Hz) survived correction (FDR corrected p < 0.05, rank-sum with 5000 shuffles across subjects z-scores). B) Aggregate normalized power for slow theta and fast theta for the anterior hippocampus (top panel) and posterior hippocampus (bottom panel). Tick marks on the top in magenta indicate time points at which normalized power for associative hits is significantly higher than misses using rank-sum with 1000 shuffles, and cyan indicates the opposite. C) Bar plots show the averaged normalized power for the time points that are significantly different across associative hits and misses. Error bars indicate standard error of the mean.

To further visualize the above described effects, we averaged oscillatory power during the time windows when significant differences were identified between associative hits and misses (Fig 2C and 3C). The averaged values demonstrate the pattern described above during retrieval, with a seemingly clear dissociation between slow and fast theta oscillations. We directly tested for this dissociation with an ANOVA model that employed the factors of memory condition (associative hit and miss), theta band (slow and fast), and hippocampal location (anterior and posterior). Consistent with the impression given by Figure 3C, the ANOVA revealed a significant interaction between memory condition and theta frequency band (F(1,208)= 16.94, p=0.0001), with great fast theta power for associative misses and greater slow theta power for associative hits. Bayesian independent samples t-test revealed that power is similar (BF_01_= 4.59) for anterior and posterior hippocampi during retrieval for slow theta oscillations.

### 5.3. Oscillatory activity in theta

To test for oscillatory activity during study and test, we employed a previously validated oscillation detection algorithm for both anterior and posterior electrode contacts (the MODAL algorithm (Watrous et al., 2018)). Plots in Figure 4 show the results for this analysis during the study (subsequent associative hits) and test (associative hits). In the anterior hippocampus during study (encoding), MODAL detected a peak in the slow theta band in 56% of electrodes and a peak in the fast theta band in 51% of electrodes; 21% of electrodes exhibited a peak in both bands simultaneously (Figure 4A). We tested for an association between slow and fast theta oscillations using Yule’s Q (Lewis-Beck et al., 2003), which revealed a moderate association between slow and fast theta oscillations in the anterior hippocampus (Q = 0.38). During test (retrieval), 40% of electrodes demonstrated an oscillation in the slow theta band, and 63% in the fast theta band in the anterior hippocampus, and 16% exhibited a peak in both bands simultaneously (Figure 4A). Yule’s Q revealed a substantial association between slow and fast theta oscillations (Q=0.63). This pattern is consistent with previous observations (Lega et al., 2012), and suggests that while anterior hippocampal electrodes may preferentially exhibit one or the other theta oscillations, both can be present simultaneously for a significant fraction of time, which may indicate that fast and slow theta oscillations reflect independent neurophysiological phenomena.

**Figure 4:**
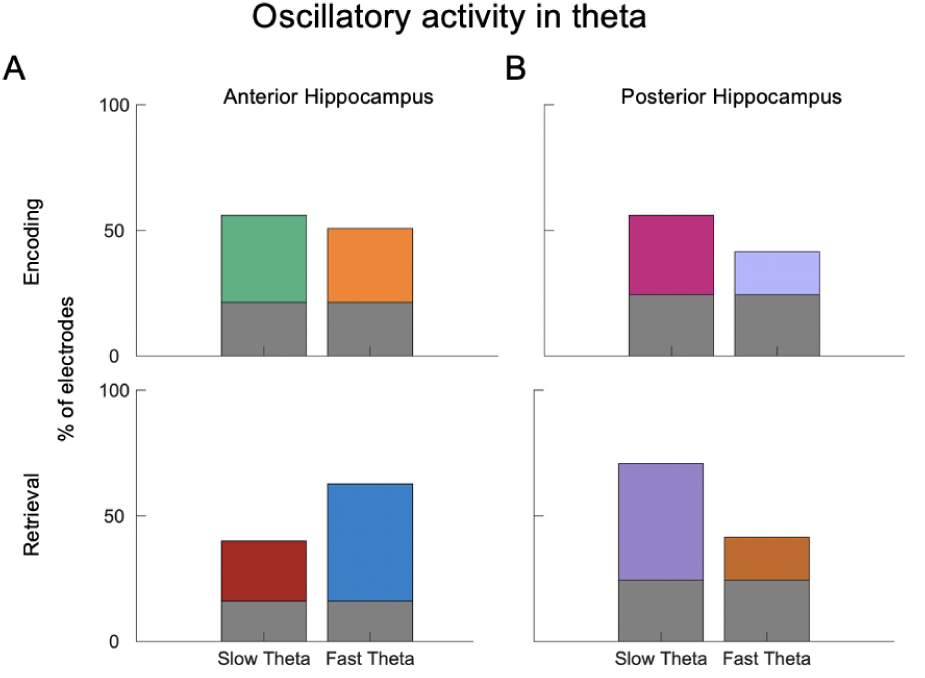
Application of multiple oscillation detection algorithm to identify peak frequencies. A bar plot showing the results of applying a peak detection algorithm to all electrodes in A) anterior hippocampus B) posterior hippocampus, in both cases during encoding (top panel) and retrieval (bottom panel) for associative hits. The height of the bar indicates the percentage of electrodes pooled across participants exhibit an oscillatory peak. Gray color bar in each plot indicates the percentage of electrodes exhibit a peak in both slow and fast theta frequency bands.

The MODAL algorithm also identified oscillations in both frequency bands in the posterior hippocampus (Figure 4B). During study, MODAL detected a peak in the slow theta band in 56% of electrodes and in the fast theta band in 41% of electrodes; 24% of electrodes exhibited a peak in both bands simultaneously. At test, 71% of electrodes demonstrated an oscillation in the slow theta band, and 41% in the fast theta band, with 24% exhibiting a peak in both bands simultaneously. There was a weak association between slow and fast theta oscillations during study (Q=-0.05) and test (Q=0.14) in the posterior hippocampus.

At the subject level, MODAL detected both slow and fast theta oscillations in 79% of individuals for the anterior hippocampus and 50% of individuals for the posterior hippocampus during study. 63% of individuals exhibited both slow and fast theta oscillations for the anterior hippocampus and 56% of individuals for the posterior hippocampus during test.

### 5.4. Phase reset supports associative memory formation

We further characterized the oscillatory power effects identified in our initial analyses by examining the phenomenon of phase reset. The goal was to determine whether phase reset distinguishes subsequent associative hits versus subsequent associative misses (recollection versus familiarity differences during encoding) and associative hits versus misses (differences during retrieval). The plots in Figure 5 show the results of this analysis. In the anterior hippocampus, study pairs elicited both slow band and fast theta band phase resets (Figure 5A) for subsequent associative hits but not subsequent associative misses (rank-sum test with shuffle procedure, FDR corrected *p* < 0.05, effect size 0.14-0.42), a phenomenon that may have contributed to the power decrease observed for the latter items during encoding (Figure 2A). During the test, associative hits elicited significantly greater phase reset than misses (rank-sum test with shuffle procedure, FDR corrected *p* < 0.05, effect size 0.14-0.42) in both the anterior and posterior hippocampus for the slow theta band (Figure 5B). While phase reset was significant across subjects (i.e. each individual phase distribution was significantly non-uniform by Rayleigh test), we did not observe a consistent and specific phase to which oscillations were resetting (no significant clustering of the mean preferred phase to which reset occurred, Figure 5C). We went on to test whether the slow theta phase reset observed for associative hits during encoding and retrieval tended to occur at a similar phase on an electrode by electrode basis. We compared the mean phase across study and test trials at the time segment of peak phase reset in a pairwise manner. The resulting distribution is shown in Figure 5D. For the anterior hippocampus, the phase difference between encoding and retrieval was significantly non–uniform, amounting to 10 degrees (*p* < 0.01, Rayleigh test), while for the posterior hippocampus the difference was not significant (*p* = 0.39). These findings suggest that greater reset of phase occurs during successful associative memory formation, and further that the angles of reset are similar between encoding and retrieval for the anterior hippocampus (mean difference approximately 10 degrees). For fast theta oscillations during encoding, we observed a distinction between earlier time points following item presentation when phase reset matched the pattern for slow theta oscillations (greater for associative hits, see Figure 5A right panel). Later time points show the opposite pattern, however, this appears to be a result of low phase reset for subsequent associative hits rather than a substantial increase for subsequent associative misses.

**Figure 5:**
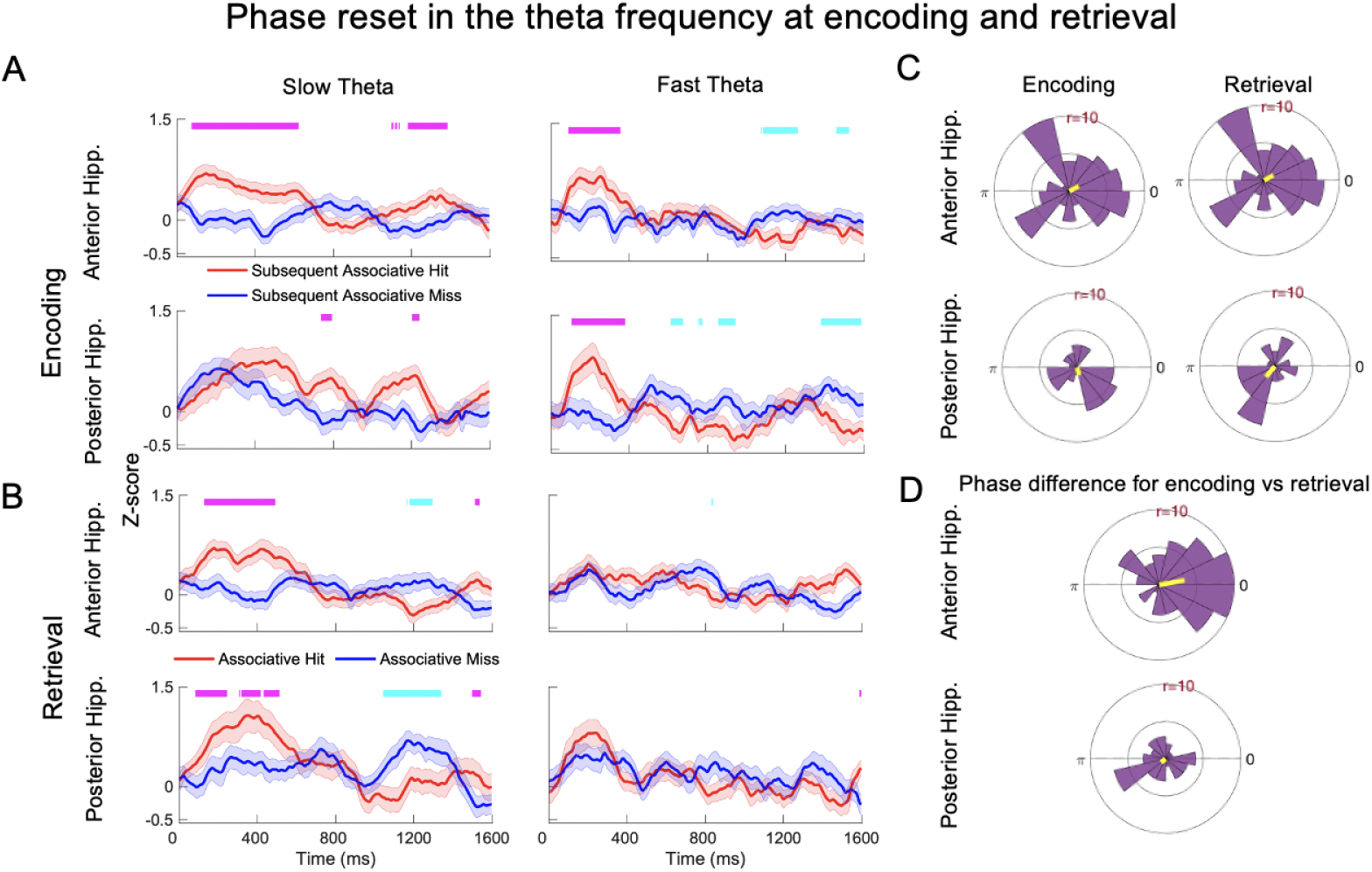
Phase reset during encoding and retrieval. Average Z-score values from Rayleigh test across all electrodes during encoding (A) and retrieval (B) are shown for the anterior hippocampus (top panel) and posterior hippocampus (bottom panel). Tick marks on the top in magenta indicate time points at which the z-score for associative hits is significantly higher than misses using rank-sum with 1000 shuffles, and cyan indicates the opposite. C) The polar histograms show the distribution of average phase values across all electrodes for slow theta in anterior (top panel) and posterior (bottom panel) hippocampus. D) Histograms of the difference in average phase values for encoding and retrieval for slow theta. The average difference is 10 degrees (non-uniform distribution) for the anterior hippocampus (top panel) and -142 degrees for the posterior hippocampus (bottom panel).

### 5.5. Reinstatement of encoding–related oscillations is stronger for recollected items

We measured the magnitude of pattern reinstatement of oscillatory activity using cosine similarity. Reinstatement quantifies the similarity pattern of brain activity elicited by a given item at encoding and retrieval. We employed the cosine similarity measure based upon its previous application to human intracranial data (Yaffe et al., 2014). We compared the magnitude of reinstatement for associative hits and misses. We focused a *priori* on reinstatement across the entire frequency spectrum from 2–200 Hz, but a breakdown for slow theta, fast theta, and high gamma (110-200 Hz) frequency bands is also shown in Figure 6. Reinstatement was significantly stronger for recollected items in both the anterior and posterior hippocampus (rank-sum test with shuffle procedure, FDR corrected *p* < 0.05, effect size 0.14-0.34), with retrieval–related activity occurring slightly earlier in time relative to encoding (above the diagonal line in Z-score plot in third column in Figure 6A and B). Reinstatement in the slow–theta range was similar in magnitude for associative hits and misses in the posterior hippocampus, but strongly differed between these classes of items in the anterior hippocampus in the latter half of the time series, i.e. approximately 1300 ms after item presentation (FDR corrected *p* < 0.05, rank-sum test). Overall, these observations are consistent with the hypothesis that reinstatement of encoding– related hippocampal activity is stronger for recollected items (associative hits) than for familiar ones.

**Figure 6:**
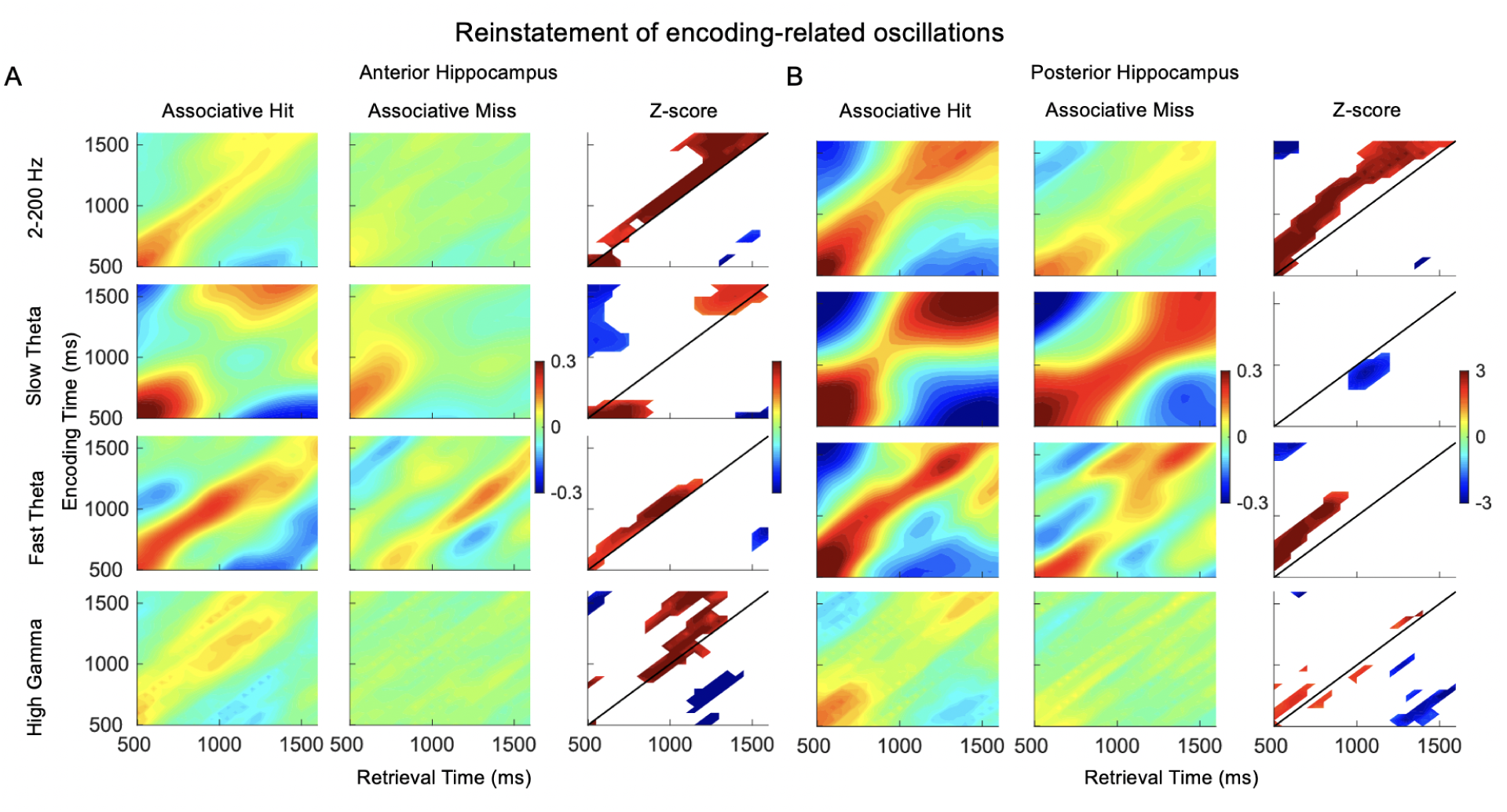
Reinstatement of encoding-related oscillations. Average cosine similarity between encoding and retrieval for associative hits and misses for A) anterior hippocampus and B) posterior hippocampus for all frequencies (2-200 Hz), slow theta, fast theta and high gamma frequency bands are shown. The plots in the third column depict encoding-retrieval pixels that survive correction. Red indicates that reinstatement for associative hits is significantly greater than associative misses, and blue indicates the opposite.

## 6. Discussion

We employed an associative recognition memory task in humans coupled with intracranial recordings. The task allowed us to directly test whether theta oscillatory activity distinguishes successful versus unsuccessful associative encoding and retrieval. This contrast allowed us to examine the key hypothesis that theta oscillations preferentially support associative recognition at the time of study (encoding) and test (retrieval). In addition, we were able to examine recollection related effects within two different theta oscillatory bands and also to compare these effects at two locations on the longitudinal axis of the hippocampus. Our key findings were that 1) power in the 2–5 Hz range reflected processes supporting recollection rather than familiarity during encoding and retrieval, 2) this power difference was driven in part by phase reset elicited by the onset of the study and test items, 3) the strength of reinstatement of oscillatory patterns distinguished recollection from familiarity based retrieval, and 4) power differences, phase reset, and reinstatement are similar between the anterior and posterior hippocampus. These observations directly link theta range activity to associative memory encoding and retrieval and establish a behaviorally–relevant contrast between 2–5 Hz slow theta and 5–9 Hz fast theta oscillations.

Models of associative memory posit that theta oscillations in the hippocampus and neocortex provide a critical binding function to link together activity across cortical areas involved in the spatially (and temporally) distributed representation of event features (Backus, Schoffelen, Szebényi, Hanslmayr, & Doeller, 2016; Staudigl & Hanslmayr, 2013). The link between theta oscillations and memory encoding in rodents has been unequivocally established through experiments demonstrating that pharmacological or surgical interruption of theta generation leads to deficits in memory formation (Fedor, Berman, Muizelaar, & Lyeth, 2010). In humans however, evidence supporting the importance of theta oscillations for memory encoding and retrieval have been somewhat mixed, as discussed in the *Introduction*. The aim of clarifying the mnemonic role of human hippocampal theta oscillations motivated us to implement the present associative recognition memory paradigm coupled with direct brain recordings: we hypothesized that if, as predicted, theta oscillations specifically support the formation of inter-item associations, then the behavioral contrast between recollection and familiarity directly operationalized by this task should elicit clear effects in the theta frequency range during both encoding and retrieval. In the same associative recognition procedure, hippocampal BOLD activity for associative hits has been reported to be greater than that for associative misses at both encoding and retrieval (de Chastelaine et al., 2016a, 2016b).

A key outstanding ambiguity regarding hippocampal theta oscillations in humans is whether there are functional differences between oscillations in the 2–5 Hz and the 5–9 Hz frequency ranges. Memory–related effects in the 2–5 Hz range have previously been reported during episodic memory encoding (Lega et al., 2012; Lin et al., 2017), spatial memory encoding and retrieval (Ekstrom et al., 2005), and recognition memory tasks (Rutishauser, Ross, Mamelak, & Schuman, 2010). However, phase locking of single unit activity in the 5–9 Hz range has been reported to differ according to rated confidence in a recognition memory paradigm (Rutishauser et al., 2015) and oscillatory differences in this band (such as phase reset and cross frequency coupling) may support working memory (Axmacher et al., 2010; Chaieb et al., 2015; Mormann et al., 2005). We separately analyzed activity in these two frequency bands seeking to elucidate whether these oscillations represent distinct phenomena by testing whether they exhibit different properties during associative memory formation and retrieval.

The most profound differences in memory–related patterns between slow and fast theta oscillations occurred during the retrieval phase, when slow theta power was significantly greater for associative hits while fast theta power was greater for associative misses (although slow theta distinguished these conditions slightly later in time following item presentation, see Figure 3B). These functional patterns support the conclusion that 2–5 Hz and 5–9 Hz oscillations in the human hippocampus are functionally distinct. The findings do not however support a model in which slow theta oscillations exclusively support associative memory formation at both encoding and retrieval: like slow oscillations, fast theta oscillations also exhibited a phase reset memory effect in both the anterior and posterior hippocampus *during encoding*. Further, the relatively greater power decrease occurring during encoding for associative misses was also similar for fast and slow theta. Understanding different connectivity relationships for these two oscillations using the associative recognition paradigm may help illuminate separate slow and fast theta networks that explicate regions involved in familiarity versus recollection, especially during retrieval. Certainly, a comprehensive understanding of these oscillations requires further characterization, including contrasts between spatial navigation and verbal memory and updated models of theta generation and propagation accounting for possible slow versus fast theta differences (Miller et al., 2018; Solomon, Lega, Sperling, & Kahana, 2019). However, our findings support a model by which slow theta oscillatory power increases are necessary for associative memory during item retrieval and as such they constitute direct evidence of a role for these oscillations in recollection.

The resetting of theta phase provides a possible mechanism for synchronizing single unit activity in the hippocampus, facilitating long term potentiation and formation of neuronal ensembles representing features of a memory event (Jutras & Buffalo, 2010). Our findings suggest that slow theta phase reset occurs during both encoding and retrieval, supporting its importance for associative encoding. The pattern for fast theta oscillations is more nuanced, as these oscillations exhibited phase reset during encoding but not retrieval for associative hits, and stronger phase reset for associative misses beginning 800 ms after item presentation. These findings echo previous human studies of *cortical* brain activity, in which recognition memory test items elicited phase reset across a broad frequency range that included fast theta oscillations (Rizzuto et al., 2006). However, temporal lobe oscillations recorded as participants performed this task exhibited different phases between encoding and retrieval, consistent with our observations for fast theta but not slow theta oscillations, which reset to a similar phase (Figure 5C). Intracranial recordings in non–human primates performing a recognition memory test also suggest that stimulus–induced reset of the phase of fast theta oscillations (centered at 8 Hz) at the time of encoding predicts retrieval success (Killian, Potter, & Buffalo, 2015). These findings reflect our observations of increased fast theta phase reset during encoding beginning approximately 800 ms after item presentation. Examining properties such as spike– field coherence separately for fast versus slow theta oscillations in this paradigm may help elucidate these differences.

Reinstatement effects were markedly stronger for associative hits than for misses in both the anterior and posterior hippocampus. This finding supports the widely held assumption that the accuracy of associative memory retrieval depends upon the re-representation of oscillatory features present at encoding (Rugg, Johnson, Park, & Uncapher, 2008). The timing of reinstatement (slightly earlier during retrieval) is consistent with previous human data obtained using a paired associates task (Yaffe, Shaikhouni, Arai, Inati, & Zaghloul, 2017). Our findings are also consistent with a prior human iEEG study that reported that memory reinstatement was identifiable in hippocampal low-frequency oscillations (1-8 Hz) during a spatial navigation task (Estefan et al., 2019). Human noninvasive data also identify reinstatement in the hippocampus stronger for recollection than familiarity–based retrieval (Liang & Preston, 2017). Additionally, our findings support models of mnemonic processing in which vivid representations characteristic of episodic memories occur during retrieval as a consequence of hippocampal pattern completion (Hanslmayr, Staresina, & Bowman, 2016; Staresina et al., 2016).

Our data do not provide strong evidence of a functional dissociation along the hippocampal longitudinal axis for either associative encoding or retrieval. The key patterns in anterior and posterior locations were similar, with theta power differences between associative hits and misses driven by power decreases for misses during encoding but power increases in the slow theta range for associative hits during retrieval. While these findings do not directly support recent models of hippocampal longitudinal specialization (such as the ‘granularity’ model proposed by Poppenk (2013)), they do suggest that both anterior and posterior hippocampus support verbal associative memory, suggesting that models focusing on emotional content (favoring anterior activation) or spatial memory (favoring posterior activation) somewhat incomplete (Poppenk et al., 2013).

